# Identification of risk variants and characterization of the polygenic architecture of disruptive behavior disorders in the context of ADHD

**DOI:** 10.1101/791160

**Authors:** Ditte Demontis, Raymond Walters, Veera M. Rajagopal, Irwin D. Waldman, Jakob Grove, Thomas D. Als, Søren Dalsgaard, Marta Ribases, Jonas Grauholm, Marie Bækvad-Hansen, Thomas Werge, Merete Nordentoft, Ole Mors, Preben Bo Mortensen, ADHD Working Group of the Psychiatric Genomics Consortium (PGC), Bru Cormand, David M. Hougaard, Benjamin M. Neale, Barbara Franke, Stephen V. Faraone, Anders D. Børglum

## Abstract

Attention-Deficit/Hyperactivity Disorder (ADHD) is a childhood psychiatric disorder often comorbid with disruptive behavior disorders (DBDs). ADHD comorbid with DBDs (ADHD+DBDs) is a complex phenotype with a risk component that can be attributed to common genetic variants. Here we report a large GWAS meta-analysis of ADHD+DBDs based on seven cohorts in total including 3,802 cases and 31,305 controls. Three genome-wide significant loci were identified on chromosomes 1, 7, and 11. A GWAS meta-analysis including a Chinese cohort supported the locus on chromosome 11 to be a strong risk locus for ADHD+DBDs across European and Chinese ancestries (rs7118422, P=3.15×10^-10^, OR=1.17). This locus was not associated with ADHD without DBDs in a secondary GWAS of 13,583 ADHD cases and 22,314 controls, suggesting that the locus is a specific risk locus for the comorbid phenotype.

We found a higher SNP heritability for ADHD+DBDs (h^2^_SNP_ =0.34) when compared to ADHD without DBDs (h^2^_SNP_ =0.20). Genetic correlations of ADHD+DBDs with aggressive (r_g_ =0.81) and anti-social behaviors (r_g_=0.82) were high, and polygenic risk score analyses revealed a significant increased burden of variants associated with ADHD and aggression in individuals with ADHD+DBDs compared to ADHD without DBDs. Our results suggests that ADHD+DBDs represent a more severe phenotype with respect to the genetic risk load than ADHD without DBDs, in line with previous studies, and that the risk load to some extent can be explained by variants associated with aggressive behavior.

## Introduction

Attention-Deficit/Hyperactivity Disorder (ADHD) is a common childhood onset behavioral disorder affecting around 5% of children and 2.5% of adults^1^. Comorbidity with other psychiatric disorders is common among children, and disruptive behavior disorders (DBDs) are the most frequently occurring conditions^2^. DBDs comprise oppositional defiant disorder and conduct disorder. Both have a childhood onset and are characterized by persistent patterns of of oppositional, defiant, disobedient and disruptive behaviour and antisocial rule-breaking and aggressive behaviours like being destructive, physically cruel towards others, and rule violations^3^. DBDs have a prevalence of 3-10% percent^4, 5^ among children and, are around twice as frequent in males compared to females^6^ and DBDs are associated with a 3-fold increased risk of premature death^7^, with higher mortality rates than in individuals with ADHD^8^. Different comorbidity rates of ADHD with DBDs have been reported, some studies have found that around 30% of children with ADHD have comorbid DBDs (ADHD+DBDs)^6, 9^, while a study of 1.92 million individuals from Denmark, found that 17% of those with ADHD were also diagnosed with DBDs^8^. Among those with DBDs in the same Danish cohort, more than half (57%) also had a comorbid diagnosis of ADHD^8^. ADHD in combination with diagnosed DBDs or excessive aggressive and disruptive behaviors increases the risk for several detrimental outcomes including increased risks for substance use disorders^10, 11^, in-patient psychiatric admission^12^, transgression^13, 14^, risky behavior^7^, and premature death compared to those only diagnosed with ADHD^7, 8^.

Both genetic and environmental factors influence the risk for ADHD as well as DBDs with twin heritability estimates of 0.74^15^ and 0.40-0.70^16–18^, respectively. Twin studies have also suggested that ADHD+DBDs is a more severe and genetically loaded subtype of ADHD than ADHD without comorbid DBDs^19^. Siblings of individuals with ADHD+DBDs have a higher recurrence risk to develop ADHD+DBDs compared to siblings of individuals having ADHD without DBDs^20, 21^. Individuals with ADHD+DBDs also have an increased polygenic burden of common ADHD risk variants compared to individuals with only ADHD, further supporting the hypothesis that ADHD+DBDs reflect a higher load of genetic risk^22^. However, it seems unlikely that ADHD risk variants alone can fully account for the underlying genetic risk that mediates aggressive and disruptive behaviours in individuals with ADHD+DBDs. Family studies have found that ADHD and DBDs have distinct genetic architectures with moderate to high genetic overlap in the range of 0.34 - 0.74^23,  24, 25^. The existence of genetic risk factors specific to the aggressive and disruptive component of ADHD+DBDs finds support from a twin study where DBDs had an estimated heritability of 0.33-0.64 after controlling for ADHD, with a significant genetic component also observed for DBDs in individuals without ADHD^26^.

Several genome-wide association studies (GWASs) have focused on diagnosed DBDs^27, 28^ or aggressive and anti-social behaviors^29, 30^, with only limited success in identifying genome-wide significant loci and no conclusive, replicated findings^27, 29, 30^. Only two genome-wide studies have focused specifically on ADHD+DBDs. One small genome-wide linkage study examined DBDs in individuals with ADHD^31^ and another, while not assessing DBD diagnoses, examined the aggressive component in individuals with ADHD^32^. Neither studies reported genome-wide significant loci.

In the current study we performed the largest GWAS-meta analysis of ADHD+DBDs to date using a Danish nation-wide cohort from iPSYCH and samples from the Psychiatric Genomics Consortium (PGC). We report the first three genome-wide significant loci for ADHD+DBDs, located on chromosomes 1, 7, and 11, and show generalization to a Chinese cohort for the locus on chromosome 11. Characterizing the polygenic architecture of ADHD+DBDs revealed a high genetic overlap with childhood aggression in the general population and anti-social behavior, considerably higher than found for ADHD without DBDs.

## Results

### GWAS meta-analysis and generalization across European and Chinese ethnicities

The meta-analysis included data from the Danish iPSYCH cohort (2,155 cases, 22,664 controls) and six European ancestry PGC cohorts (1,647 cases, 8,641 controls). All cases were diagnosed with both ADHD and DBDs or had a diagnosis of hyperkinetic conductict disorder which according to the ICD10 criteria implies that both phenotypes are present. Selection of controls was population-based and they were not diagnosed with ADHD or DBDs. Results were in total based on 3,802 cases and 31,305 controls and included 8,285,688 variants after filtering. Three loci passed the threshold for genome-wide significance (P=5×10^-8^); these were located on chromosome 1 (index variant rs549845, P=2.38×10^-8^, OR=1.16), 7 (index variant rs11982272, P=4.38×10^-8^, OR=0.83), and 11 (index variant, rs7118422, P=8.97×10^-9^, OR=1.16) (Table 1, Figure 1A and Supplementary Figure 1.A-C). The directions of association of the index variants in the three loci were consistent across all cohorts (Supplementary Figure 2A-C).

**Figure 1.**
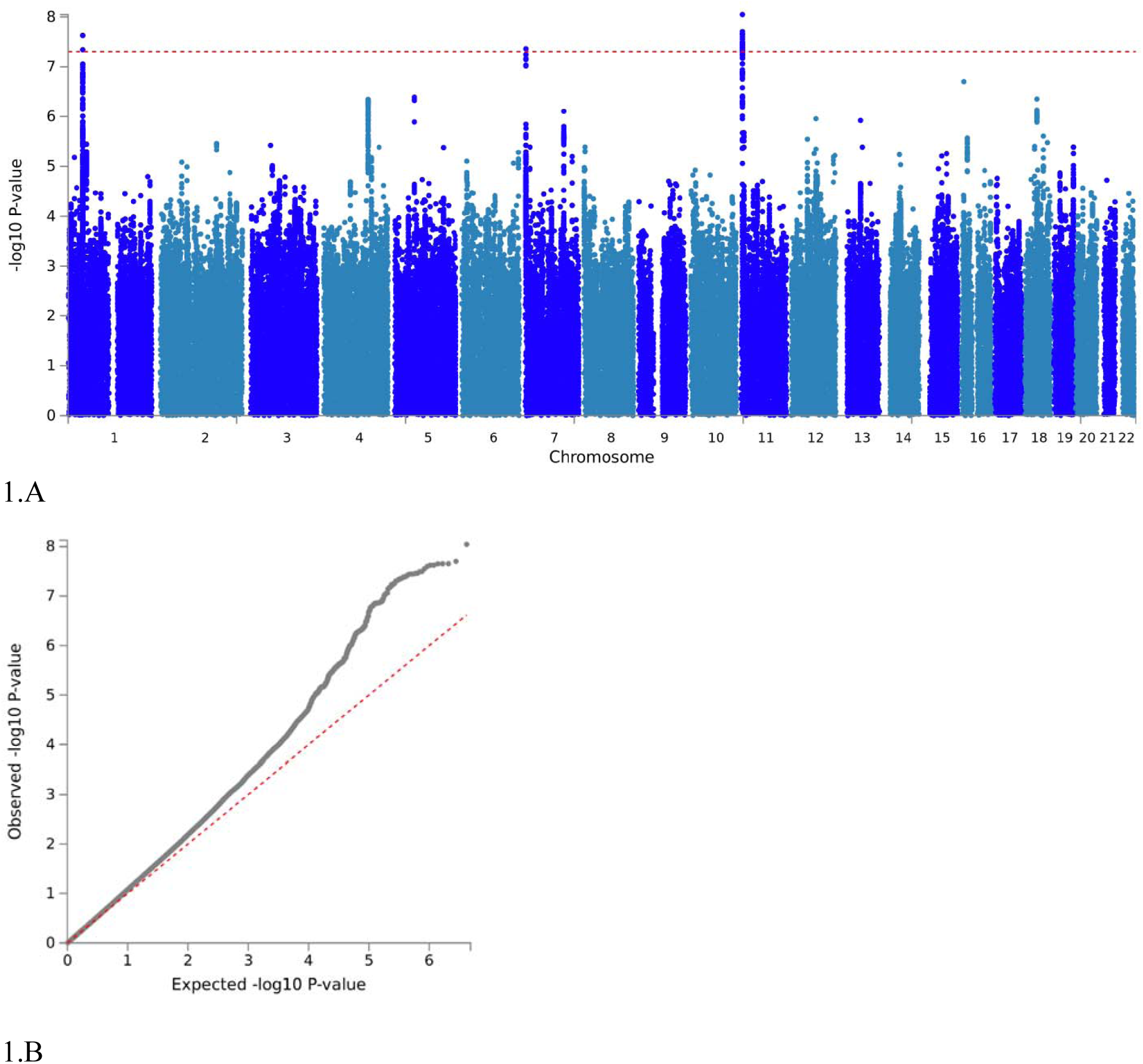
Manhattan plot and q-q plot of results from the GWAS meta-analysis of ADHD+DBDs. **A.** Results from GWAS meta-analysis of iPSYCH and PGC cohorts in total including 3,802 cases and 31,305 controls. Two-sided P-values from meta-analysis using an inverse-variance weighted fixed effects model. The red vertical line represents the threshold for genome-wide significant association (P = 5×10^-8^). **B.** Quantile-quantile plot of the -log10 P-values from the GWAS meta-analyses. The dotted line indicates the distribution under the null hypothesis.

**Table 1.** Results for the genome-wide significant index variants in the three loci associated with ADHD+DBDs identified in the GWAS meta-analysis of 3,802 individuals with ADHD+DBDs and 31,305 controls (iPSYCH+PGC). Results from the trans-ancestry GWAS meta-analysis for identification of cross-ethnicity risk variants in Europeans and Chinese (iPSYCH+PGC+Chinese). The location (chromosome (chr)), gene location of index variant (Gene), alleles (A1 and A2), odds ratio (OR) of the effect with respect to A1, and association P-values from inverse-variance weighted fixed effects model of the index variants are given.

| SNP | Chr | Gene | A1 | A2 | iPSYCH+PGC |  |  | Chinese |  |  | iPSYCH+PGC+Chinese |  |  |
| --- | --- | --- | --- | --- | --- | --- | --- | --- | --- | --- | --- | --- | --- |
|  |  |  |  |  | 3,802 cases |  |  | 406 cases |  |  | 4,208 cases, |  |  |
|  |  |  |  |  | 31,305 controls |  |  | 917 controls |  |  | 32,222 controls |  |  |
|  |  |  |  |  | OR | SE | P | OR | SE | P | OR | SE | P |
| rs7118422 | 11 | *STIM1* | T | C | 1.164 | 0.026 | $8.97 \times 10^{-9}$ | 1.276 | 0.089 | 0.006 | 1.173 | 0.025 | $3.15 \times 10^{-10}$ |
| rs549845 | 1 | *PTPRF* | G | A | 1.166 | 0.027 | $2.38 \times 10^{-8}$ | 0.943 | 0.095 | 0.541 | 1.147 | 0.026 | $2.179 \times 10^{-7}$ |
| rs11982272 | 7 | *MAD1L1* | T | C | 1.199 | 0.033 | $4.39 \times 10^{-8}$ | 0.901 | 0.132 | 0.132 | 1.179 | 0.032 | $3.013 \times 10^{-7}$ |

### Homogeneity of effects in the PGC and iPSYCH cohorts and intercept evaluation

To evaluate the consistency of the genetic architecture underlying ADHD+DBDs in iPSYCH and the PGC cohorts, we estimated the genetic correlation between the two using LD score regression^33, 34^. The genetic correlation between the iPSYCH cohort and the meta-analyzed PGC cohorts and was high (r_g_=0.934, SE=0.14, P=3.26×10^-11^) supporting consistency of the ADHD+DBDs phenotypes analyzed in the cohorts. In addition, no variants demonstrated significant heterogeneity between studies (Supplementary Figure 3 and 4).

LD Score regression analysis indicated that the observed deviation of the genome-wide test statistics from the null distribution (lambda=1.11, Figure 1B) was mainly caused by polygenicity. The intercept ratio estimate suggests that the majority of the inflation of the mean J statistic of the GWAS meta-analysis is attributable to polygenic effects (ratio=0.87, SE=0.0662) rather than confounding factors. The estimated remaining contribution of confounding factors was small and non-significant (intercept =1.015; SE=0.008; P=0.064).

### Trans-ancetry GWAS meta-analysis

To replicate and generalize the findings to other ethnicities, a GWAS of ADHD+DBDs was performed in a Chinese cohort (406 cases, 917 controls) and a fixed effects meta-analysis including the Chinese, European iPSYCH and PGC cohorts was performed (total 4,208 cases and 32,222 controls). The three loci identified in the main analysis were evaluated for significance in the trans-ancestry GWAS meta-analysis. The locus on chromosome 11 was nominally significant in the Chinese cohort (P=0.006), and the association P-value in the meta-analysis became stronger in the trans-ancestry GWAS meta-analysis (P=3.15×10^-10^, OR=1.17) (Figure 2 and Supplementary Figure 5) suggesting the chromosome 11 locus to be a risk locus for ADHD+DBDs across ethnicities. The results incorporating the Chinese cohort did not support replication of the other two loci (Table 1).

**Figure 2.**
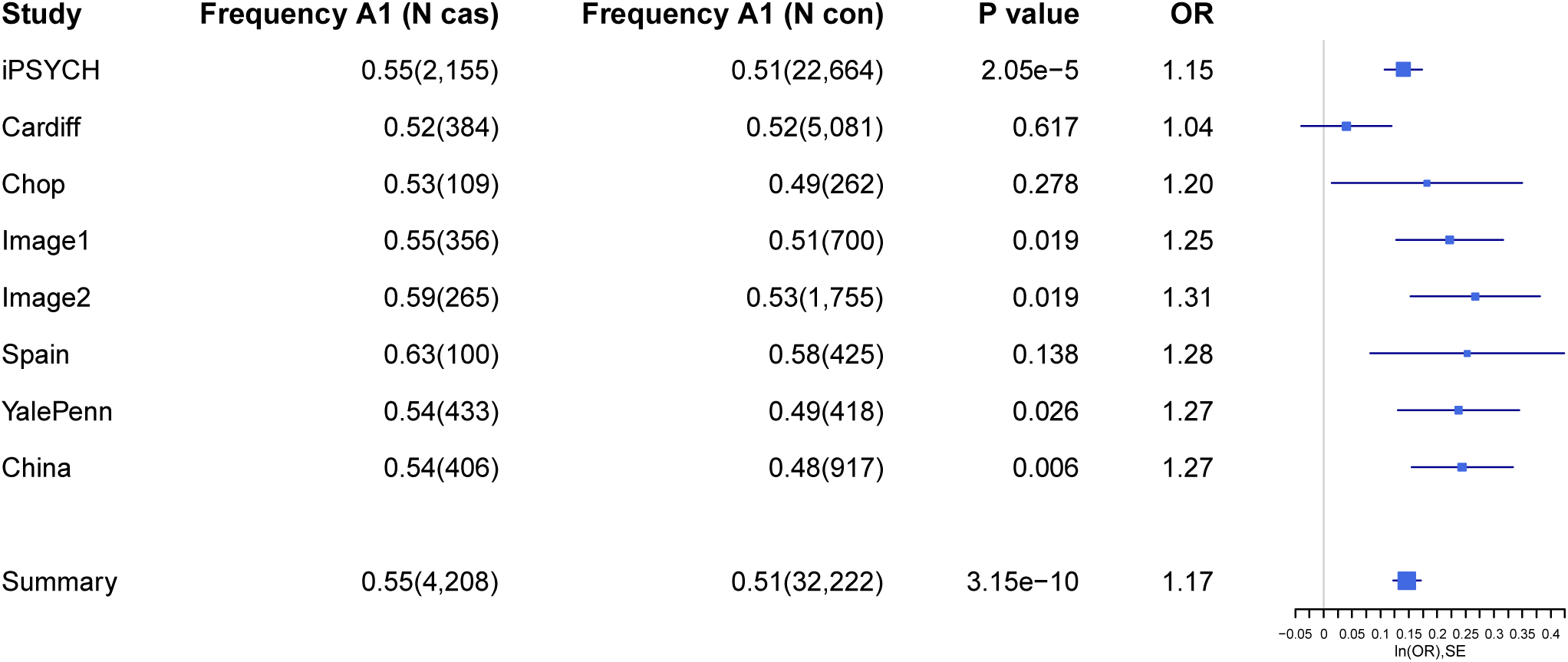
Forest plot for index variant in the genome-wide significant locus for ADHD+DBDs. Forest plot for the index variant (rs7118422) in the genome-wide significant locus on chromosome 11. Results from the generalization GWAS meta-analysis across European and Chinese ancestries including 4,208 cases and 32,222 controls. Freqeuncy of the risk variant (Frequency A1) in cases and controls in the included cohorts are given together with sample size in cases (N cas) and controls (N con). P-values and odds ratios (OR) from the GWASs. The plot provides a visualization of the effect size estimates (natural logarithm of the odds ratio (ln(OR)) in each included cohort, estimated from logistic regression, and for the summary meta-analysis using an inverse-variance weighted fixed effects model. In addition, standard errors of the ln(OR) estimates.

### Secondary GWASs

For subsequent evaluation of how much of the signal in the GWAS meta-analysis of ADHD+DBDs was driven by the oppositional/aggressive component of the comorbid phenotype, two additional GWASs were conducted using iPSYCH samples. To adjust for the effect of ADHD, we performed a case-only GWAS comparing 1,959 individuals with ADHD+DBDs against 13,539 individuals having ADHD without DBDs, referred to as the “ADHD+DBD vs. ADHD-only GWAS”. Additionally, a GWAS of 13,583 cases having ADHD without DBDs and 22,314 population-based controls, referred to as the “ADHD without DBDs GWAS”, was performed (the case-control numbers differ between GWASs due to deviation in the numbers of related individuals and genetic outliers removed in the analyses).

The summary statistics from the two secondary GWASs were used to evaluate the direction of association of the top loci (281 loci, P< 1=10^-4^) from the GWAS meta-analysis of ADHD+DBDs. A consistent direction of association was observed for 221 loci (out of the 281 loci) in the ADHD without DBDs GWAS (sign test P < 2.2=10^-16^), while all the 281 loci demonstrated consistent direction of association in the ADHD+DBD vs. ADHD-only GWAS. The proportion of variants having a consistent direction of association in the ADHD+DBD vs. ADHD-only GWAS was significantly larger than the proportion in the ADHD without DBDs GWAS (P = 7.7=10^-16^), which suggests that the associations in the GWAS meta-analysis of ADHD+DBDs reflects association with the comorbid phenotype beyond association with risk for ADHD alone.

Of the significant loci from the GWAS of ADHD+DBDs, the loci on chromosomes 7 and 11 had nominally stronger effect sizes in the ADHD+DBD vs. ADHD-only GWAS compared to the ADHD without DBDs GWAS, while the chromosome 1 locus had similar effects in both secondary GWASs (Supplementary Table 9). The difference is most striking for rs7118422 on chromosome 11, which showed minimal evidence for association with ADHD without DBDs (OR=1.022, p=0.175) versus more strongly suggestive evidence of association with ADHD+DBD vs. ADHD-only (OR=1.126, p=7.07 =10^-04^). To help formalize this comparison, we used mtCOJO^35^ to estimate the joint effects of the significant loci from the GWAS of ADHD+DBDs conditional on effects mediated through the genetics of ADHD without DBDs. Under this model (see Methods) none of the three loci reached genome-wide significance for a direct effect on ADHD+DBDs, although the locus on chromosome 11 retained the most robust signal after correction for ADHD without DBDs (OR_adjusted_=1.14; P_adjusted_=1.43=10^-06^).

### Gene-based association test

A gene-based association analysis was performed using MAGMA^36^. Six genes (*RRM1*, *STIM1, MAML3*, *ST3GAL3*, *KDM4A* and *PTPRF*) were significantly associated with ADHD+DBDs (P < 2.7=10^-6^; correcting for 18,553 genes analyzed; Supplementary Figure 6 and Supplementary Table 4). Three genes (*ST3GAL3*, *KDM4A* and *PTPRF)* are located in the genome-wide significant locus on chromosome 1, and two genes *(RRM1*, *STIM1)* in the genome-wide significant locus on chromosome 11. One gene *(MAML3)* was located on chromosome 4 and had not been identified as a risk locus in the single variant analysis.

To evaluate if the gene-based association signals reflected the aggressive and disruptive component of the ADHD+DBDs phenotype rather than ADHD alone, we did a gene-set test of the most associated genes from the primary ADHD+DBDs GWAS meta-analysis (P<10^-3^, 79 genes) using the results from the two secondary GWASs. The gene set was significantly associated with ADHD without DBDs (beta=0.312(SE=0.02), P=9=10^-4^), but had a much stronger association in the ADHD+DBD vs. ADHD only GWAS (beta=1.1 (SE=0.07), P = 9.28=10^-32^).

### Association of genetically regulated gene expression with ADHD+DBDs

Association of genetically regulated gene expression with ADHD+DBDs was analyzed in 12 brain tissues from GTEx^37^ (version 6p) using MetaXcan^38^. Depending on the tissue, 2,042-6,094 genes were tested (Supplementary Table 4). Three genes were differently expressed in ADHD+DBDs cases compared with controls after Bonferroni correction (correcting for the total number of tests performed (43,142); P<1,16=10^-6^); *RRM1* (chromosome 11) was less expressed in cases, and *RAB3C* (chromosome 5) and *LEPRE1* (chromosome 1) showed a higher expression in cases when compared to controls (Supplementary Table 4). The genes on chromosome 1 and 11 were located in or near genome-wide significant loci, whereas the locus on chromosome was novel.

### SNP heritability

The estimated h^2^_SNP_ for ADHD+DBDs was 0.25 (SE=0.03) using LD score regression^33^ with a prevalence estimate of 2% in the population (Supplementary Table 6). When considering only the iPSYCH cohort, a higher SNP heritability was found for ADHD+DBDs (h^2^_SNP_ =0.34; SE=0.05) compared to the GWAS of ADHD without DBDs (h^2^_SNP_ =0.20; SE=0.02). This pattern remained similar when using GCTA and was stable when assuming a lower prevalence of ADHD+DBDs and higher prevalence of ADHD without DBDs (Supplementary Table 6). Additionally, common variants explained a small fraction of the variation in the ADHD+DBDs phenotype compared to the ADHD without DBDs phenotype (h^2^_SNP_ =0.08; SE=0.04, GCTA estimate) (Supplementary Table 6).

### Genetic correlation with aggression related phenotypes

We estimated the genetic correlations of the ADHD+DBDs phenotype with aggression related phenotypes using GWAS results from analyses of aggressive behaviors in 18,988 children^29^ (EAGLE aggression) and antisocial behavior in 16,400 individuals^30^ (Broad Antisocial Behaviour Consortium (BroadABC)) using LD score regression^33^. We found a strong genetic correlation of ADHD+DBDs with aggression in children (EAGLE aggression, r_g_=0.81, P=0.001) and anti-social behavior (BroadABC, r_g_=0.82, P=0.007) (Supplementary Table 7). In contrast, ADHD without DBDs was only significantly correlated with aggression in children (r_g_=0.74, P=4.6=10^-5^). Analyzing only the iPSYCH cohort, ADHD+DBDs demonstrated a nominally higher positive genetic correlation than ADHD without DBDs with aggression in children (r_g_=0.85, P=5=10^-4^; and r_g_=0.74, P=4.58=10^-5^, respectively) and anti-social behavior (r_g_= 0.92, P=9=10^-3^; and r_g_=0.56, P=0.01, respectively). The differences in the genetic correlations, however, were not statistically significant when assessed using the jackknife method.

### ADHD+DBD vs. ADHD-only polygenic risk score analysis

Case-only polygenic risk score (PRS) analyses were done to evaluate whether ADHD+DBDs cases are enriched for variants associated with 22 relevant phenotypes compared with cases having ADHD without DBDs. Investigating PRS from phenotypes related to personality, cognition, and psychiatric disorders, seven phenotypes were significantly associated with ADHD+DBDs compared with ADHD without DBDs after multiple testing correction (Supplementary Table 8). Significantly increased PRS for aggressive behavior^29^ (Z=4.80, P=1.51=10^-6^, OR=1.13) and ADHD (Z=5.42, P=5.90=10^-8^, OR=1.21) were observed. Additionally, PRS for increased cognitive performance was negatively associated with ADHD+DBDs compared to ADHD without DBDs (educational attainment^39^, Z=-3.30, 8.0=10^-4^, OR=0.92; college or university degree^40^ Z=-3.22, P=1.0=10^-3^, OR=0.93; human intelligence^41^ Z=-3.00, P=2.00=10^-3^, OR=0.93; verbal-numerical reasoning^40^ Z=-3.26, P=1.00=10^-3^, OR=0.92). Finally, PRS for having children at a younger age was associated with ADHD+DBDs compared to ADHD without DBDs (Z=-4.40, P=8.4=10^-6^, OR=0.9). Only a small proportion of the variance in ADHD+DBDs (compared to ADHD without DBDs) was explained by the PRSs, with the maximum Nagelkerke’s *R^2^* = 0.36% for the ADHD PRS. The odds ratio for ADHD+DBDs was found to increase with increasing polygenic load of variants associated with aggression, ADHD, and age at first birth, and to decrease with higher load of variants associated with cognition (Supplementary Figures 7. A-G). The highest risk was observed for the ADHD PRS, where the 20% of ADHD cases with the highest ADHD PRS had an OR=2.48 for having comorbid DBDs relative to the 20% with the lowest ADHD PRS (Supplementary Figure 7.A).

## Discussion

This study is the first to identify genome-wide significant loci for ADHD+DBDs based on a meta-analysis of 3,802 cases and 31,305 controls from the iPSYCH cohort and six cohorts from PGC. We identified three risk loci on chromosomes 1, 7, and 11 with odds ratios ranging from 1.16 to 1.20, in line with what was found in the recent GWAS meta-analysis of ADHD^42^. These risk loci demonstrated high consistency in the direction of association in the included cohorts, indicating that the associations likely have a true biological cause rather than being spurious signals driven by one or few cohorts (Figure 2 and Supplementary Figure 2.A-C). The high genetic correlation observed between the PGC cohorts and the iPSYCH cohort suggests that the genetic architecture underlying ADHD+DBDs were similar in the two samples. In the GWAS meta-analysis for generalization of the identified risk loci to a Chinese sample, only the locus on chromosome 11 replicated the findings in the European samples. This locus seems to be specifically associated with the aggressive and disruptive component of the ADHD+DBDs phenotype, since the effect remained strong when comparing comorbid ADHD+DBD to ADHD alone but disappeared in the GWAS of ADHD without DBDs (Supplementary Table 9).; consistent with this, evidence for a direct effect of the locus on ADHD+DBDs remained after adjusting for the effect of ADHD alone in the mtCOJO analysis (Supplementary Table 3).

In contrast, the locus on chromosome 1, which had been identified as a strong risk locus for ADHD previously^42^, seems to reflect an association with ADHD. This locus remained genome-wide significant in the GWAS of ADHD without DBDs (Supplementary Table 9) and the association with ADHD+DBDs decreased considerably in the analyses adjusting for the effect of ADHD without DBDs and in the ADHD+DBD vs. ADHD-only GWAS (Supplementary Table 3 and 9).

The locus on chromosome 7 seems to be a shared risk locus between ADHD+DBDs and ADHD without DBDs. The locus remained moderately associated in the GWASs adjusted for ADHD (Supplementary Table 3 and 9) as well as in the ADHD without DBDs GWAS (Supplementary Table 9). The locus is located in *MAD1L1*, which is involved in mitotic spindle-assembly checking before anaphase. The locus is novel with respect to ADHD and DBDs, but was found genome-wide significant in the recent large cross-disorder GWAS^43^ and has previously been associated with schizophrenia and bipolar disorder^44, 45, 46^ suggesting that *MAD1L1* could be a cross-disorder risk gene for several psychiatric disorders.

The locus most strongly associated with ADHD+DBDs on chromosome 11 is located in *STIM1* (Supplementary Figure 1C), a gene not previously implicated in ADHD, DBDs, aggression-related phenotypes, or psychiatric disorders. *STIM1* encodes a transmembrane protein (STIM1) in the endoplasmatic reticulum (ER) that acts as a sensor of calcium. Upon calcium depletion from the ER, STIM1 is responsible for an influx of calcium ions from the extracellular space through store-operated calcium channels to refill ER stores^47–49^. Store-operated calcium entry may also be involved in neuronal calcium signaling^50^, and recent evidence indicates that STIM1 plays a role in synaptic plasticity affecting learning and memory^50, 51^. These results are interesting in the light of the observed learning deficits associated with aggressive behaviors and accumulating evidence that suggests calcium signaling to be involved in several psychiatric disorders^52–54^. Alternatively, analysis of genetically regulated gene expression suggested that the variants in the genome-wide significant locus might affect expression of *RRM1*, with a decreased *RRM1* expression being associated with ADHD+DBDs. *RRM1* is oriented in a tail-to-head configuration with *STIM1*, which lies 1.6 kb apart, and is involved in the biosynthesis of deoxyribonucleotides from the corresponding ribonucleotides necessary for DNA replication. To our knowledge, this gene has not previously been associated with psychiatric disorders.

Six genes were exome-wide significantly associated with ADHD+DBDs, including two implicated by variants in or near the genome-wide significant locus on chromosome 11 (*RRM1* and *STIM1*) and with three (*ST3GAL3*, *KDM4A*, and *PTPRF*) out of the remaining four located in or near the genome-wide significant locus on chromosome 1. The top-associated genes (79 genes) seem to mainly to reflect association with the aggressive and disruptive component of the ADHD+DBDs phenotype, since they were more strongly associated in the ADHD+DBD vs. ADHD-only GWAS (where the effect of ADHD is corrected out) than in the ADHD without DBDs GWAS. Likewise, the most strongly associated single markers (with P<1=10^-4^) in the GWAS meta-analysis of ADHD+DBDs showed higher consistency in the direction of association in the ADHD+DBD vs. ADHD-only GWAS than in the GWAS of ADHD without DBDs, reinforcing the idea that the associations mainly reflect the aggressive and disruptive component of the phenotype.

When evaluating the polygenic architecture of ADHD+DBDs, a higher SNP heritability was found for ADHD+DBDs (h^2^_SNP_ =0.34) compared to ADHD without DBDs (h^2^_SNP_ =0.2) (Supplementary Table 5). These estimates are consistent with the recently reported SNP heritability of ADHD (h^2^_SNP_ =0.22)^42^, which included individuals with and without comorbid DBDs. Conditional on an ADHD diagnosis, the aggressive and disruptive behavioral component of the ADHD+DBDs phenotype also has a genetic component involving common variants (h^2^_SNP_ =0.08). The SNP heritability of ADHD+DBDs compared to ADHD without DBDs is partly explained by a higher burden of common risk variants for ADHD, supported by the PRS analysis (Supplementary Table 8). The significantly higher burden of ADHD risk variants among individuals with ADHD+DBDs was especially evident when examining individuals belonging to the 20% of ADHD cases with the highest ADHD genetic risk load, who had an odds ratio of 2.48 for having comorbid DBDs (Supplementary Figure 7A). This is also a replication of previous findings of a higher load of ADHD risk variants in individuals with ADHD and comorbid conduct disorder compared to those having only ADHD^22^.

Going beyond ADHD risk burden, the common variant component of DBDs could also include variants distinctly associated with aggression. This idea is supported by our finding of increased PRS for aggression in ADHD+DBDs compared to ADHD without DBDs (Supplementary Figure 7B). This conclusion is reinforced by the genetic correlation results, where we found somewhat higher genetic correlation of ADHD+DBDs with both aggressive behavior in children^29^ and anti-social behavior^30^ (Supplementary Table 7) compared to those found for ADHD without DBDs. Additionally, these results imply that the genetic architecture underlying the aggressive and disruptive behavioral component of the ADHD+DBDs phenotype overlaps strongly with that affecting aggressive and anti-social behavior in the general population. Thus, aggressive and anti-social behaviors seem to have a continuous distribution in the population, with individuals having ADHD+DBDs representing an extreme. This is in line with what has been observed for other complex phenotypes, such as diagnosed ADHD representing the upper tail of impulsive and inattention behaviors^42^, and diagnosed autism spectrum disorder representing the upper tail for social communication difficulties and rigidity^55, 56^.

Aggressive behavior is stable across age intervals during childhood^57^, and twin studies have suggested genetics to play an important role in this stability^57, 58^. Moreover, early aggression might be predictive of later serious anti-social behaviour^59^ resulting in increased risk of getting a diagnosis of antisocial personality disorder^60^. Our results suggest that common genetic variants play an important role in childhood aggression, which has also been reported previously^29^, and that the subsequent risk for anti-social behavior in individuals with ADHD+DBDs to some extent has an underlying biological cause involving common variants.

We identified individuals with ADHD+DBDs to have increased load of variants associated with worse cognition compared to individuals having ADHD without DBDs (Supplementary Table 8; Supplementary Figure 7C-F). This could reflect the increased genetic load of ADHD risk variants in ADHD+DBDs, since ADHD has a strong negative genetic correlation with cognition-related phenotypes^42^. However, it might also involve variants associated with anti-social behavior, which also has a negative genetic correlation with educational performance^30^. This latter idea is supported by epidemiological studies linking aggression to decreased educational attainment^61–63^. Finally, we found a significantly higher load of variants associated with younger age at birth of first child in ADHD+DBDs compared to ADHD without DBDs, in line with the observed positive genetic correlations of ADHD and anti-social behavior with having children earlier^30, 42^ and evolutionary theories suggesting that aggression has played a role when competing for access to mates^64^.

In summary, we identified three genome-wide significant loci for ADHD+DBDs. The locus on chromosome 11 was associated most strongly with the comorbid phenotype, and seems to be a cross-ancestry risk locus in European and Chinese. Our results suggest that the aggressive and disruptive behavioral component of the ADHD+DBDs phenotype has a biological cause, which in part can be explained by common risk variants associated with ADHD, aggressive and anti-social behavior. Individuals with ADHD+DBDs therefore represent a more severe phenotype with respect to the genetic risk load than ADHD without DBDs, most likely including loci which are distinct to ADHD+DBDs. This study represents the first step towards a better understanding of the biological mechanism underlying ADHD+DBDs.

## Methods

### Samples – the iPSYCH cohort

The iPSYCH cohort is a population-based nation-wide cohort which includes 79,492 genotyped individuals (∼50,000 diagnosed with major psychiatric disorders and ∼30,000 controls). The cohort was selected, based on register information from a baseline birth cohort of all singletons born in Denmark between May 1st, 1981 and December 31, 2005 (N=1,472,762) (see a detailed description in^65^). A biological sample of the included individuals were obtained from the Newborn Screening Biobank at Statens Serum Institute, Denmark. DNA was extracted from dried blood spot samples and whole genome amplified in triplicates as described previously^66, 67^. Genotyping and calling of genotypes were performed as described in ^65, 68^.

For this study cases and controls were identified based on diagnoses given in 2016 or earlier in the Danish Psychiatric Central Research Register register^69^. Cases with ADHD+DBDs had a diagnosis of hyperkinetic conduct disorder (F90.1) or an ADHD diagnosis (ICD-10 F90.0) occurring together with a diagnosis of ODD (ICD-10 F91.3) or conduct disorder (ICD-10 F91.1, F91.2, F91.9, F91.9). Distribution of cases with ADHD+DBDs over diagnosis codes is presented in Supplementary Table 1. ADHD cases without DBDs were defined as individuals having ADHD (ICD-10 F90.0) without any diagnosis of DBDs. Controls were randomly selected from the same nation-wide birth cohort and not diagnosed with ADHD or DBDs.

The study was approved by the Danish Data Protection Agency and the Scientific Ethics Committee in Denmark. All analyses of the iPSYCH sample were performed at the secured national GenomeDK high performance-computing cluster (https://genome.au.dk).

### Samples - cohorts from the Psychiatric Genomics Consortium

For the meta-analysis, seven ADHD cohorts (six cohorts of European ancestry and one of Chinese ancestry) aggregated by PGC with information about diagnoses of ADHD+DBDs were included. An overview of the cohorts including genotyping information and diagnosis criteria can be found in Supplementary Table 2. Detailed descriptions of the cohorts can be found elsewhere^68^. Details on approval authorities can be found in Supplementary Table 2.

### Quality control and imputation

Quality control, imputation and primary GWASs of the iPSYCH and PGC cohorts were done separately for each using the bioinformatics pipeline Ricopili^70^. Pre-imputation quality control allowed an inclusion of individuals with a call rate > 0.98 (>0.95 for iPSYCH) and genotypes with a call rate >0.98, difference in SNP missingness between cases and controls < 0.02, no strong deviation from Hardy-Weinberg equilibrium (*P* >1=10^−6^ in controls or *P* >1=10^−10^ in cases) and low individual heterozygosity rates (| F_het_ | <0.2). Genotypes were phased and imputed using SHAPEIT^71^ and IMPUTE2^72^ and the 1000 Genomes Project phase 3 (1KGP3)^73, 74^ as imputation reference panel. Trio imputation was done with a case-pseudocontrol setup.

Relatedness and population stratification were evaluated using a set of high-quality genotyped markers (minor allele frequency (MAF) >0.05, HWE P >1=10^-4^ and SNP call rate >0.98) pruned for linkage disequilibrium (LD) resulting in ∼30,000 pruned variants (variants located in long-range LD regions defined by Price et al.^75^ were excluded). Genetic relatedness was estimated using PLINK v1.9^76, 77^ to identify first and second-degree relatives (π̂ >0.2) and one individual was excluded from each related pair (cases preferred kept over controls). Genetic outliers were excluded based on principal component analyses (PCA) using EIGENSOFT^78, 79^. For iPSYCH a genetic homogenous sample was defined based on a subsample of individuals being Danes for three generations as described in Demontis and Walters et al.^68^. For the PGC samples genetic outliers were removed based on visual inspection of the first six PCs. For all cohorts PCA was redone after exclusion of genetic outliers.

### GWAS meta-analysis and generalization across European and Chinese ethnicities

Association analysis was done using additive logistic regression and the imputed marker dosages, covariates from principal component analyses after removal of genetic outliers and other relevant covariates (Supplementary Table 2). Meta-analysis of the iPSYCH cohort (2,155 cases, 22,664 controls) and the six PGC cohorts (1,647 cases, 8,641 controls) was done using an inverse standard error weighted fixed effects model and the software METAL^80^ and included in total 3,802 cases and 31,305 controls.

For generalization of genetic risk variants across European and Chinese ancestries, a GWAS meta-analysis was done as described above including the iPSYCH cohort, the six European PGC cohorts and the cohort of Chinese ancestry. In total, 4,208 cases and 32,222 controls were included. No individual genotypes were used for the meta-analysis.

### Homogeneity of effects in the PGC and iPSYCH cohorts and intercept evaluation

LD score regression^33, 34^ was used to estimate the genetic correlation using summary statistics from GWAS of ADHD+DBDs in the iPSYCH cohort and meta-analysis of the six European PGC cohorts. Only variants with an imputation info score > 0.9 were included. The intercept was restricted to one as there was no sample overlap and no indication of population stratification.

The ratio (ratio = (intercept-1)/(mean chi^2^− 1)) from LD score regression was used to evaluate the relative contribution of polygenic effects and confounding factors to the observed deviation from the null in the genome-wide distribution of the X^2^ statistics of the GWAS meta-analysis of ADHD+DBDs.

### Secondary GWASs

In order to correct out the effect of ADHD we did a case-only GWAS comparing 1,959 individuals with ADHD+DBDs against 13,539 individuals having ADHD without DBDs, referred to as the “ADHD+DBD vs. ADHD-only GWAS”. Additionally, a GWAS of 13,583 cases having ADHD without DBDs and 22,314 population-based controls referred to as the “ADHD without DBDs GWAS” was performed. Both GWASs were based only on iPSYCH samples and performed using additive logistic regression and the imputed marker dosages, covariates from principal component analyses after removal of genetic outliers and covariates indicating the 23 genotyping waves.

The summary statistics from the two secondary GWASs were used to evaluate direction of association of the top loci associated with ADHD+DBDs using a sign test based on LD distinct variants (r^2^<0.2, 281 variants) with association P-values less than 1=10^-4^ in the GWAS meta-analysis of ADHD+DBDs.

We also did an mtCOJO^35^ analysis to estimate the effect of the top loci for ADHD+DBDs conditional on genetic effects on ADHD alone. This was done using summary statistics from the GWAS meta-analysis of ADHD+DBDs and from the GWAS of ADHD without DBDs. The analysis was run using mtCOJO^35^ implemented in GCTA^81^ using standard procedures. Following default settings, estimation of the effect of ADHD without DBDs on ADHD+DBDs (as part of the indirect path contributing to marginal ADHD+DBDs associations) was performed using variants that were genome-wide significant in the GWAS of ADHD without DBDs (P<5=10^-8^) and not in linkage disequilibrium (r^2^< 0.05; 7 index variants). No variants were removed due to evidence of pleiotropy (HEIDI-outlier threshold of P=0.01).

### Gene-based association test

Gene-based association analysis was done using MAGMA 1.05^36^ and summary statistics from the GWAS meta-analysis. Variants were annotated to genes using the NCBI37.3 gene definitions and no window around genes was used. MAGMA summarizes association signals observed for variants located in a gene into a single P-value while correcting for LD in a reference genome. For this the European samples from the 1000 Genomes phase 3 were used.

The most associated genes in the GWAS meta-analysis of ADHD+DBDs (80 genes, P<10^-3^) were evaluated in a gene-set test for association with ADHD+DBDs compared to ADHD without DBDs and for association with ADHD without DBDs. Gene-based P-values were generated using summary statistics from the two secondary GWASs (ADHD+DBDs vs ADHD-only GWAS and ADHD without DBDs GWAS) and subsequently gene-set tests were done using MAGMA 1.05^36^. MAGMA performs a competitive test to analyze if the gene set is more strongly associated with the phenotype than other genes, while correcting for a series of confounding effects such as gene length and size of the gene set.

### Association of the genetically regulated gene expression with ADHD+DBDs

Association of the genetically regulated gene expression with ADHD+DBDs was analyzed in 12 brain tissues from GTEx^37^ (version 6p) using MetaXcan^38^. MetaXcan is an extension of PrediXcan^82^ that can be used to test for differences in gene expression using summary statistics. We used high-performance prediction models for MetaXcan based on variants located within 1 Mb +/- of transcription start site and trained using elastic net regression and 10-fold cross-validation^4^ downloaded from http://predictdb.org. MetaXcan also requires covariance matrices of the variants within each gene model for each tissue. Covariance matrices calculated from 503 individuals with European ancestry from the 1000 genomes project^74^ available with the prediction models at http://predictdb.org were used.

### SNP heritability

The SNP heritability (h^2^_SNP_) was estimated using LD score regression^33^ and the summary statistics from the GWAS meta-analysis of ADHD+DBDs. The heritability was estimated on the liability scale assuming a population prevalence of ADHD+DBDs of 2% and 1%.

In order to evaluate the extent to which common genetic variants contributes to the risk of ADHD+DBDs compared to having ADHD without DBDs, the SNP heritability of for the two phenotypes were estimated only in iPSYCH samples. This was done using LD score regression and univariate GREML analyses in GCTA^81^. h^2^_SNP_ was estimated on the liability scale assuming a population prevalence of 2% and 1% for ADHD+DBDs and 3% and 4% for ADHD without DBDs. The GCTA analyses were corrected for the same covariates as used in the GWASs.

Additionally, we evaluated how much of the variance in the ADHD+DBDs phenotype could be explained by common genetic variation in the context of ADHD. For this we did a case-only approach including 1,959 cases with ADHD+DBDs and 13,539 individuals having ADHD without DBDs. This was only done using GCTA due to a low polygenic signal in the ADHD+DBD vs. ADHD-only GWAS GWAS (mean X^2^= 1.06).

### Genetic correlation with aggression related phenotypes

The genetic overlap of ADHD+DBDs with aggression related phenotypes was evaluated by estimating genetic correlations using LD score regression^33^ and the summary statistics from the GWAS meta-analysis of ADHD+DBDs and results from two aggression related GWASs. One is a GWAS meta-analysis of scores of aggressive behaviors in 18,988 children^29^ (EAGLE aggression) obtained by questionnaires filled by their parents. Another is a GWAS meta-analysis of antisocial behavior conducted by the Broad Antisocial Behaviour Consortium (BroadABC) including 16,400 individuals^30^. Both children and adults were accessed for a broad range of antisocial measures, including aggressive and non-aggressive domains. The BroadABC study has a minor overlap with the EAGLE aggression GWAS with respect to the included cohorts.

Statistical difference between two r_g_ estimates was calculated using the block jackknife method implemented in the LDSC software^33, 83^. The variants acrosse the genome were divided in 200 blocks and jackknife delete values were calculated by excluding one block at a time. The computed jackknife delete values were then used to calculate corresponding jackknife pseudo values. By using the mean and variance of the jackknife pseudovalues, Z score and corresponding P values were computed, testing the null hypothesis that the difference between the r_g_s is equal to zero.

### Polygenic risk score analysis

Case-only polygenic risk score (PRS) analyses were done using GWAS summary statistics from 22 GWASs related to cognition and education (six phenotypes), personality (nine phenotypes), psychiatric disorders (five phenotypes), reproduction/fitness (two phenotypes) (detailed list of phenotypes see Supplementary Table 8). Variants with imputation info score < 0.9, MAF < 0.01, missing values, ambiguous and multi-allelic variants, indels and duplicated identifiers were removed. The remaining variants were LD-clumped using Plink^76^. PRS was estimated at different P-value thresholds in the 22 training datasets: P<5=10^-8^, 1=10^-6^, 1=10^-4^, 1=10^-3^, 0.01, 0.05, 0.1, 0.2, 0.5 and 1.0. PRS in the target individuals (iPSYCH samples: 1,959 ADHD+DBDs cases and 13,539 ADHD without DBDs cases) were estimated multiplying the natural log of the odds ratio of each variant by the allele-dosage of each variant and whole-genome PRS was obtained by summing values over variants for each individual. The ADHD PRS was generated using the approach described in Demontis et al.^68^. For each P-value threshold the variance in the ADHD+DBDs phenotype explained by PRS was estimated using Nagelkerke’s *R* (R package ‘BaylorEdPsych’), and association of PRS with ADHD+DBDs compared to ADHD without DBDs was estimated using logistic regression including the same covariates used in the GWAS.

Subsequently individuals were divided into quintiles based on their PRS. OR for ADHD+DBDs compared to ADHD without DBDs was estimated within each quintile with reference to the lowest risk quintile (using the training data P-value threshold resulting in the highest Nagelkerke’s *R^2^* in the target data).

## Acknowledgements

The iPSYCH team was supported by grants from the Lundbeck Foundation (R165-2013-15320, R102-A9118, R155-2014-1724 and R248-2017-2003) and the universities and university hospitals of Aarhus and Copenhagen. The Danish National Biobank resource was supported by the Novo Nordisk Foundation. Data handling and analysis on the GenomeDK HPC facility was supported by NIMH (1U01MH109514-01 to ADB). High-performance computer capacity for handling and statistical analysis of iPSYCH data on the GenomeDK HPC facility was provided by the Center for Genomics and Personalized Medicine and the Centre for Integrative Sequencing, iSEQ, Aarhus University, Denmark (grant to ADB).

This project has received funding from the European Union’s Seventh Framework Programme for research, technological development and demonstration under grant agreement no 602805 (Aggressotype) and 728018 (Eat2beNICE). This work reflects only the authors’ views, and the European Union is not liable for any use that may be made of the information contained herein. Barbara Franke was supported by a personal Vici grant of the Dutch Organisation for Scientific Research (grant 016-130-669) and by a grant for the Dutch National Science Agenda for the NWA NeurolabNL project (grant 400 17 602). Raymond Walters was supported by the Stanley Center for Psychiatric Research and the National Institute of Mental Health (5U01MH109539).

Bru Cormand received support from the Spanish ‘Ministerio de Economía y Competitividad’ [grant numbers SAF2015-68341-R, RTI2018-100968-B-I00], from AGAUR, ‘Generalitat de Catalunya’ [grant number 2017-SGR-738] and from the European Union FP7 [grant agreement n° 602805] and H2020 Programs [grant agreements n° 667302, 728018]. This work was also funded by Instituto de Salud Carlos III (PI15/01789, PI16/01505, PI17/00289, PI18/01788), and co-financed by the European Regional Development Fund (ERDF), Agència de Gestió d’Ajuts Universitaris i de Recerca-AGAUR, Generalitat de Catalunya, Spain (2017SGR1461), the Health Research and Innovation Strategy Plan (PERIS SLT006/17/287), Generalitat de Catalunya, Spain, the European College of Neuropsychopharmacology (ECNP network: ‘ADHD across the lifespan’), Departament de Salut, Generalitat de Catalunya, Spain, and a NARSAD Young Investigator Grant from the Brain & Behavior Research Foundation. The research leading to these results has also received funding from the European Union Seventh Framework Program (FP72007-2013) under grant agreement No 602805 (Agressotype) and from the European Union H2020 Programme (H2020/2014-2020) under grant agreements Nos. 667302 (CoCA) and 728018 (Eat2BeNICE). Marta RiBasés is a recipient of a Miguel de Servet contract from the Instituto de Salud Carlos III, Spain (CP09/00119 and CPII15/00023). Dr. Dalsgaard’s research is supported by grants from The Lundbeck Foundation (iPSYCH grant no R102-A9118, R155-2014-1724 and R248-2017-2003), National Institute of Health (R01, grant no ES026993), Novo Nordisk Foundation (grant no 22018), the European Commission (Horizon 2020, grant no 667302), Tryg Foundation (109399) and Helsefonden (grant no 19-8-0260).

## Author contributions

This section will be filled during the revision process

