## Supplementary info for "Identification of risk variants and characterization of the polygenic architecture of disruptive behavior disorders in the context of ADHD"

### **Supplementary Figure 1A-C. Regional association plots of the genome-wide significant loci**

Regional association plots of the local association results from the three genome-wide significant loci identified in the GWAS meta-analysis of ADHD+DBDs (3,802 cases and 31,305 controls). The y-axis represents –log(P-values) of variant association; the P-values are two-sided from meta-analysis using an inverse-variance weighted fixed effects model. The vertical green line represents the threshold for genome-wide significance (P =5x10^-8^). Location and orientation of the genes in the region is indicated, LD estimates of surrounding SNPs with the index SNP (r^2^ values estimated based on 1KGP3) is indicated by colour (colour bar in upper left corner indicates r2 values. Additionally, the local estimation of recombination rate is indicated in light blue (legend on vertical axis at right). Detailed SNP info in upper right corner (blue letters): SNP name (rsid), P-value (p), odds ratio (or), minor allele frequency(maf), imputation INFO score (info), directions in the analyzed cohorts (risk increasing - decreasing - missing).

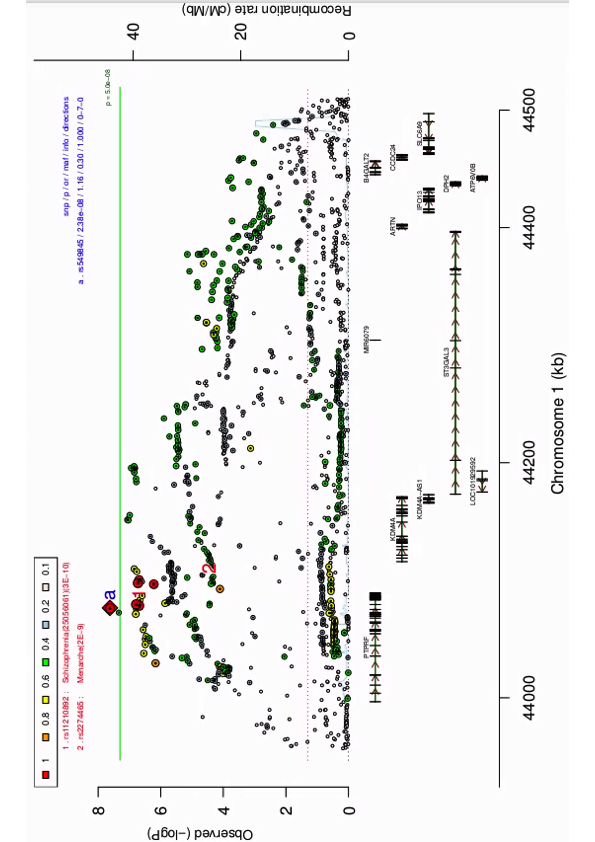

**Supplementary Figure 1A.** Regional association of the variants located the genome-wide significant locus on chromosome 1 identified in the GWAS meta-analysis of ADHD+DBDs (3,802 cases and 31,305 controls).

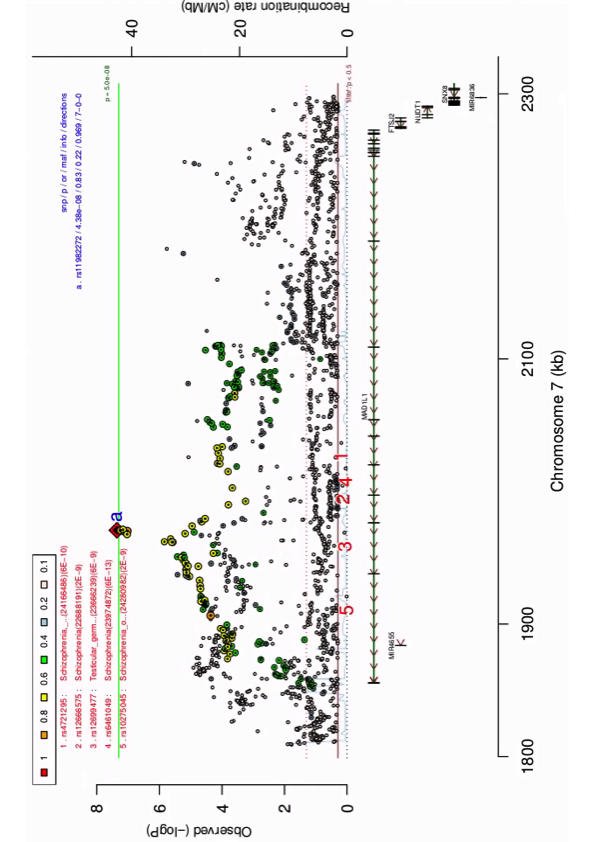

**Supplementary Figure 1B.** Regional association of the variants located the genome-wide significant locus on chromosome 7 identified in the GWAS meta-analysis of ADHD+DBDs (3,802 cases and 31,305 controls).

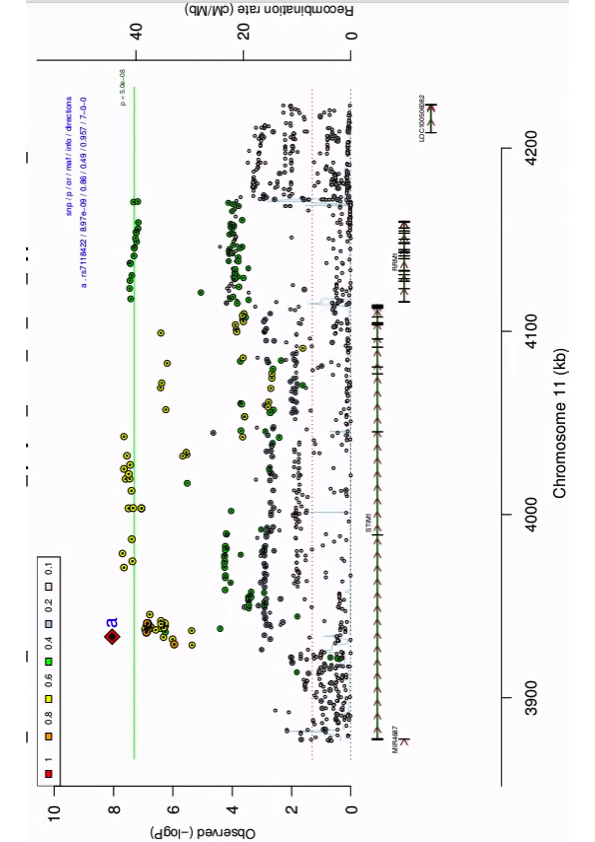

**Supplementary Figure 1C.** Regional association of the variants located the genome-wide significant locus on chromosome 11 identified in the GWAS meta-analysis of ADHD+DBDs (3,802 cases and 31,305 controls).

### **Supplementary Figure 2A-C. Forest plots of genome-wide significant index variants**

Forest plots for the index variants in the three genome-wide significant loci identified in the GWAS meta-analysis of ADHD+DBDs (3,802 cases and 31,305 controls). The plots provides a visualization of the effect size estimates (natural logarithm of the odds ratio (ln(OR)) in each included cohort, estimated from logistic regression and for the summary meta-analysis using an inverse-variance weighted fixed effects model. In addition, the standard error intervals for the effect size estimates.

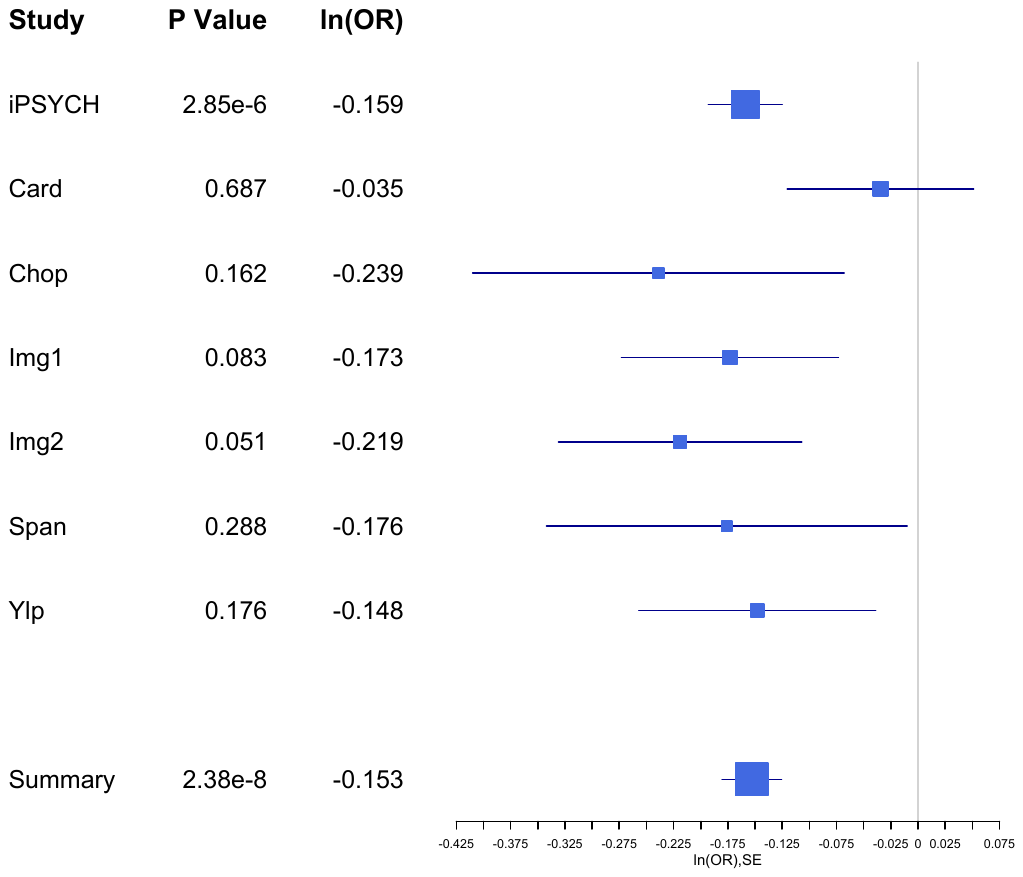

**Supplementary Figure 2A.** Forest plots for the index variant rs549845 in the genome-wide significant locus on chromosome 1 identified in the GWAS meta-analysis of ADHD+DBDs (3,802 cases and 31,305 controls). Vertical lines represent standard errors.

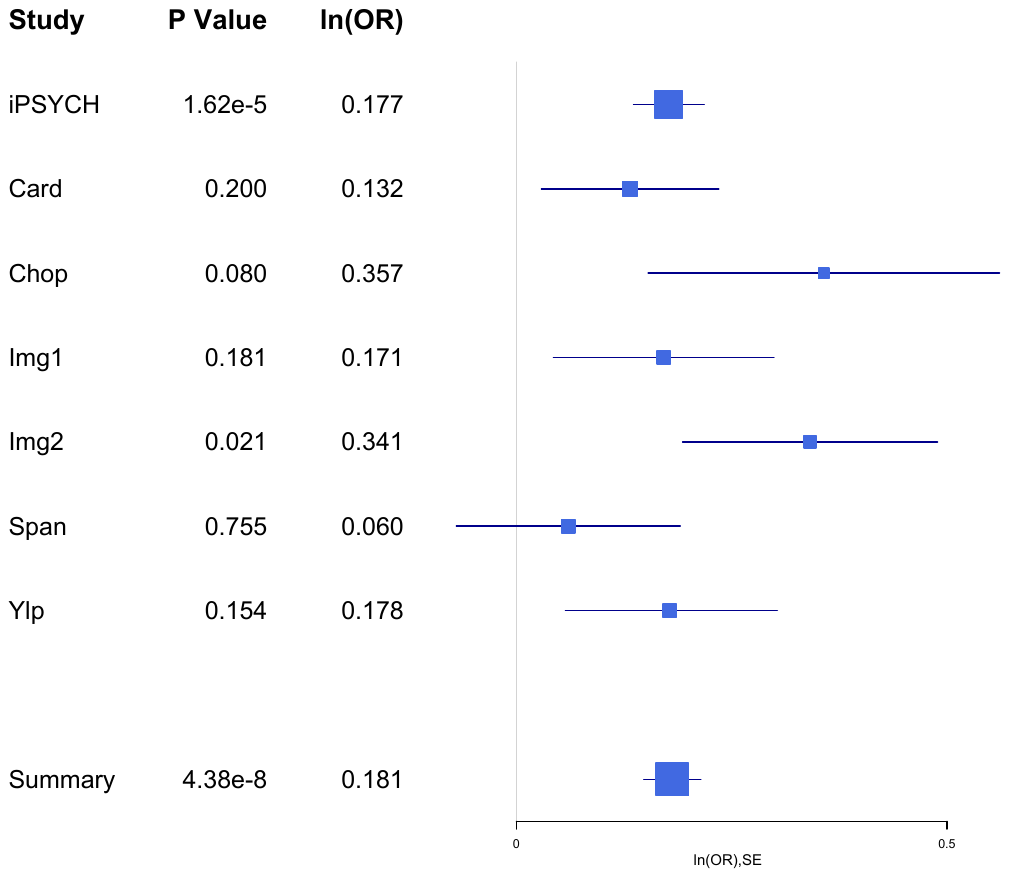

**Supplementary Figure 2B.** Forest plots for the index variant rs11982272 in the genome-wide significant locus on chromosome 7 identified in the GWAS meta-analysis of ADHD+DBDs (3,802 cases and 31,305 controls). Vertical lines represent standard errors.

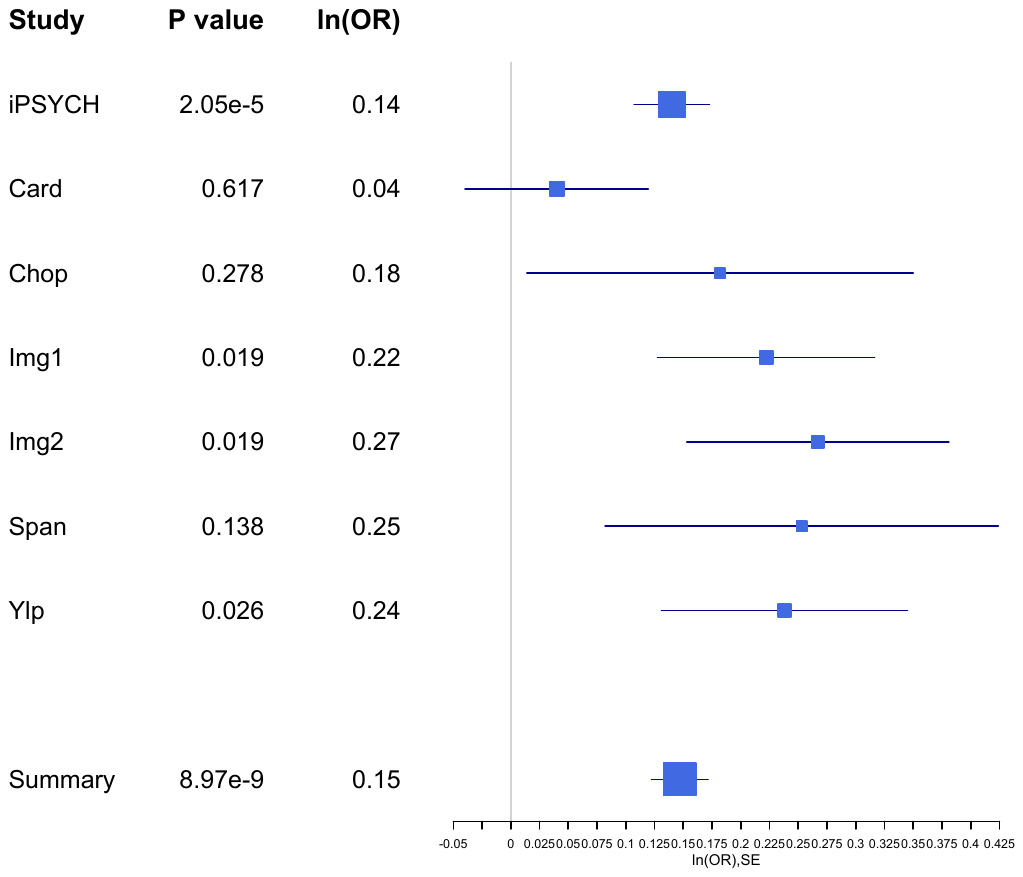

**Supplementary Figure 2C.** Forest plots for the index variant rs7118422 in the genome-wide significant locus on chromosome 11 identified in the GWAS meta-analysis of ADHD+DBDs (3,802 cases and 31,305 controls). Vertical lines represent standard errors.

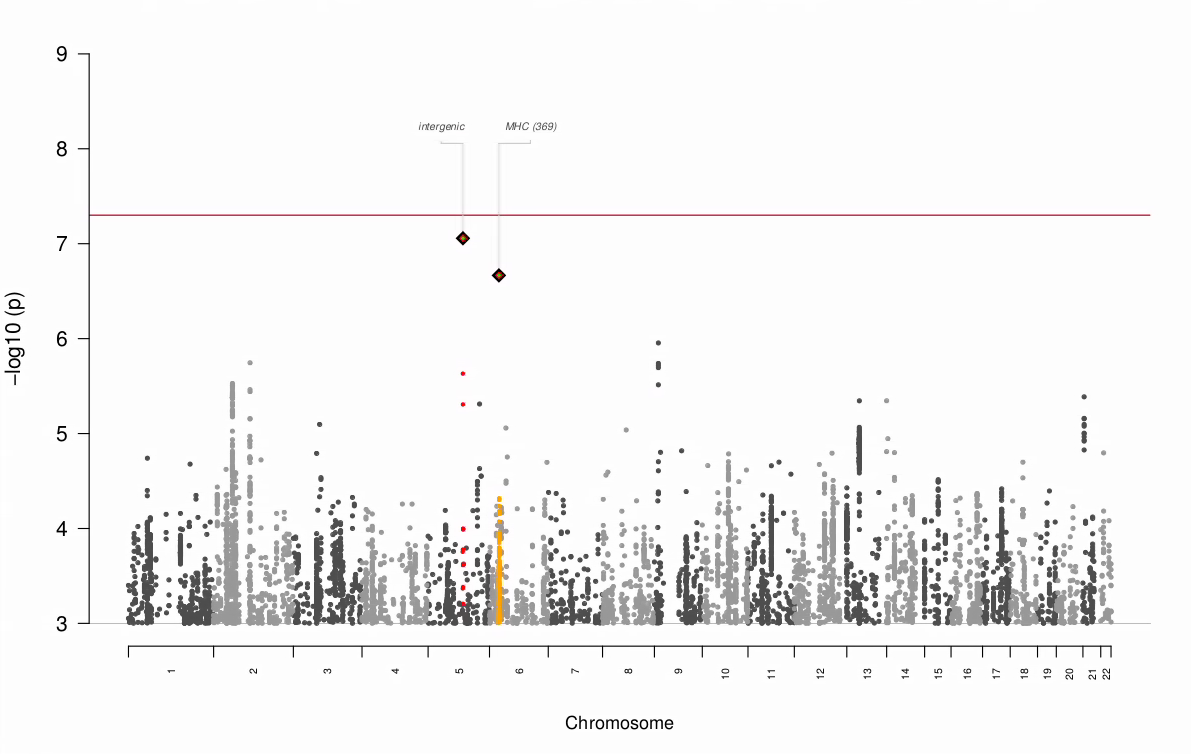

### **Supplementary Figure 3. Test for heterogeneity across cohorts**

The y-axis represents –log(P-values) from omnibus test of heterogeneity across cohorts tested with Cochran’s Q test and quantified with the I^2^ heterogeneity index. See Supplementary Table 2 for sample sizes. Red reference line indicates genome-wide significance threshold (P =5x10^-8^).

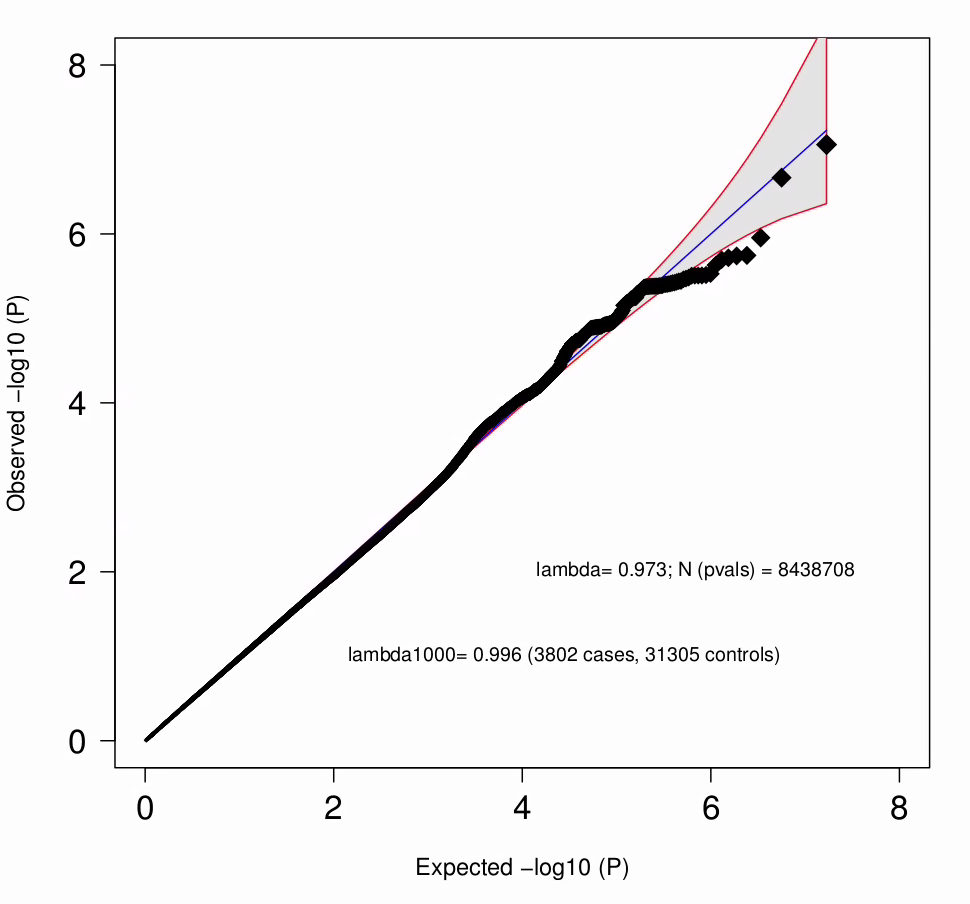

### **Supplementary Figure 4. Q-Q plot from test for heterogeneity between all cohorts in GWAS meta-analysis**

Quantile-quantile plot of P-values from the omnibus test of heterogeneity (I squared statistic (I2)) between cohorts. See Supplementary Table 2 for sample sizes of cohorts. The blue line indicates the distribution under the null hypothesis and the shaded area indicates the 95% confidence band.

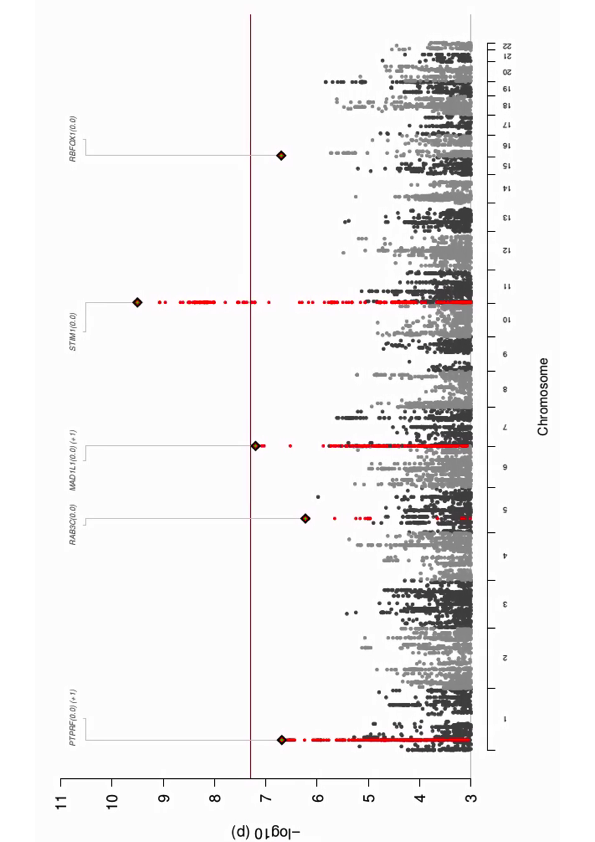

### **Supplementary Figure 5. Manhattan plot of GWAS meta-analysis of European and Chinese cohorts**

Results from cross-ancestry GWAS meta-analysis of iPSYCH and PGC cohorts with European and Chinese ancestries. Two-sided P-values from meta-analysis using an inverse-variance weighted fixed effects model, and a sample size of 4,208 cases and 32,222 controls. The red vertical line represents the threshold for genome-wide significant association (P = 5x10^-8^).

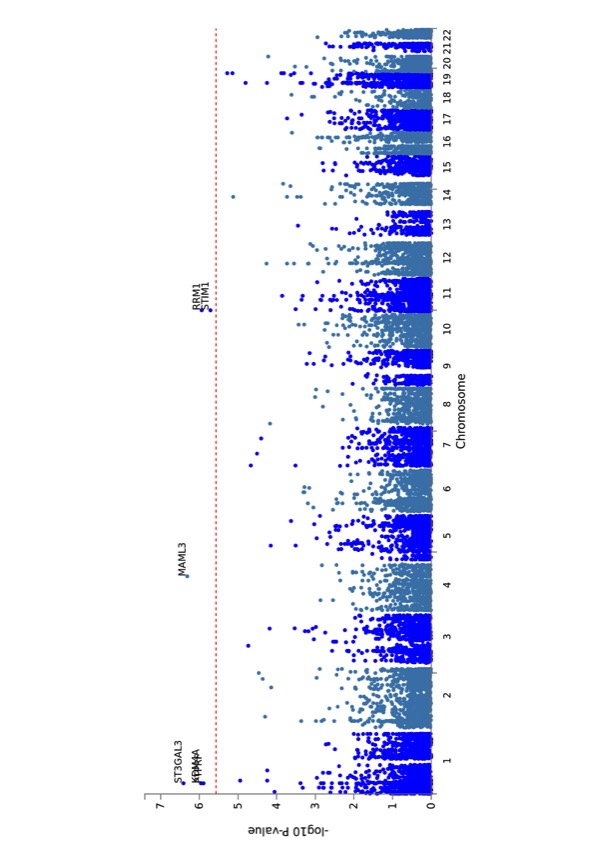

### **Supplementary Figure 6. Manhattan plot from gene-based MAGMA association analysis**

The y-axis represents –log(P-values) of gene-based association with ADHD; P-values are two-sided from MAGMA analysis based on summary statistics from the GWAS meta-analysis of 3,802 cases and 31,305 controls. The vertical red dotted line represents the threshold for exome-wide significance (P = 2.7x10^-6^).

### **Supplementary Figure 7A-G. Quintile plots of odds ratio for ADHD+DBDs by PRS**

Odds Ratio (OR) by PRS within each quintile for ADHD+DBDs compared to ADHD without DBDs. The plots represent the seven phenotypes (detailed results on Supplementary Table 8) demonstrating significant association of PRS with ADHD+DBDs after correcting for multiple testing. Error bars indicate 95% confidence limits.

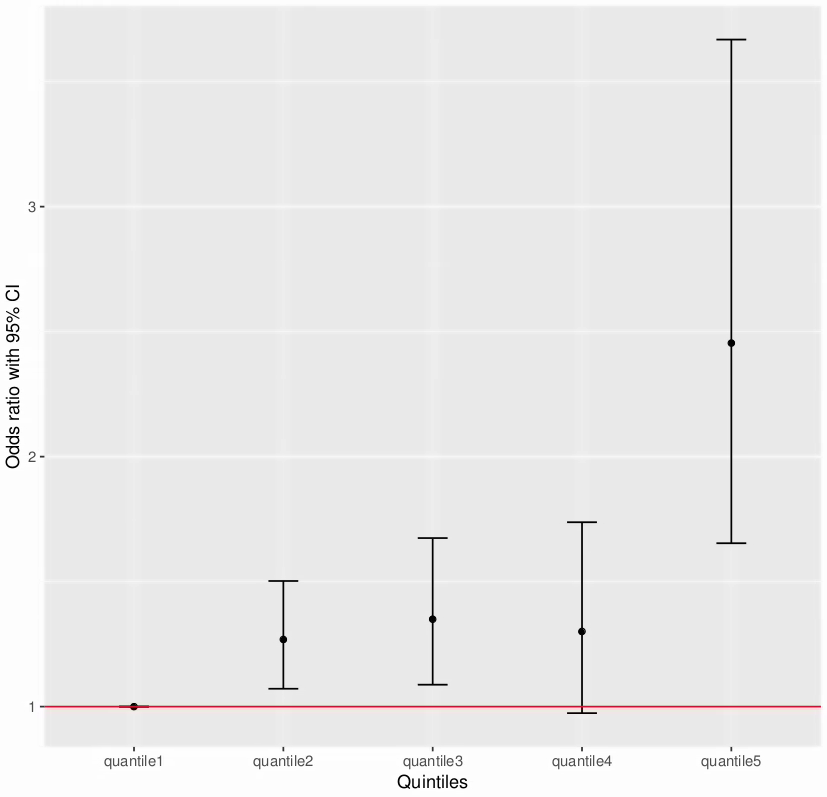

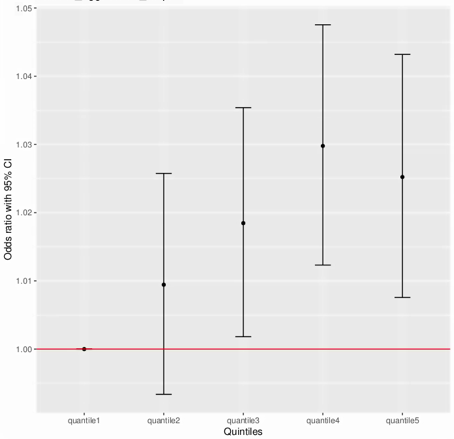

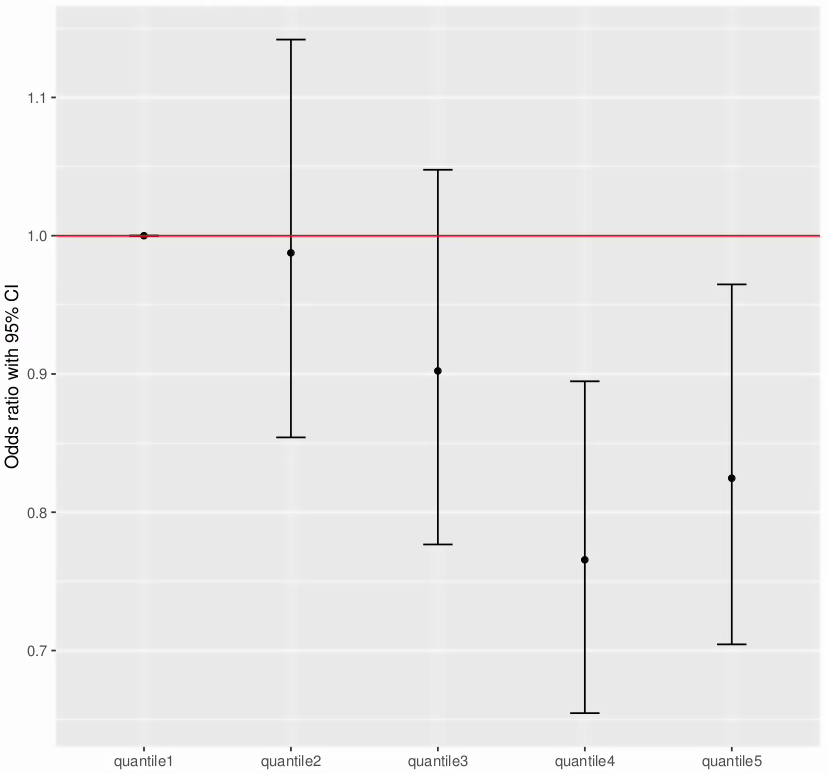

C. OR for ADHD+DBDs compared to ADHD without DBDs by PRS for number of educational years.

B. OR for ADHD+DBDs compared to ADHD without DBDs by PRS for aggression in children.

A. OR for ADHD+DBDs compared to ADHD without DBDs by PRS for ADHD.

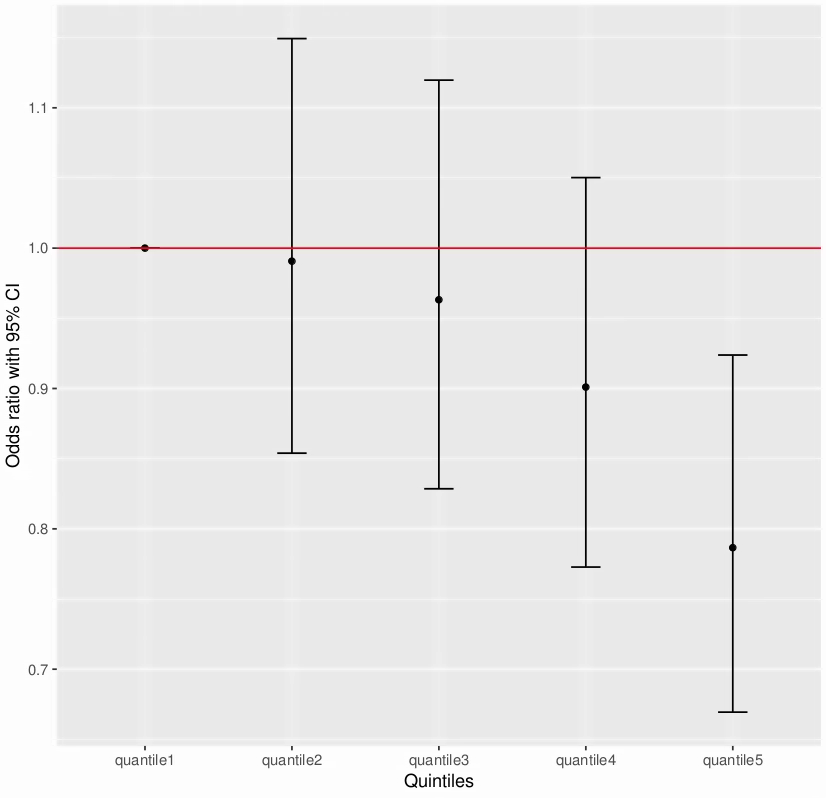

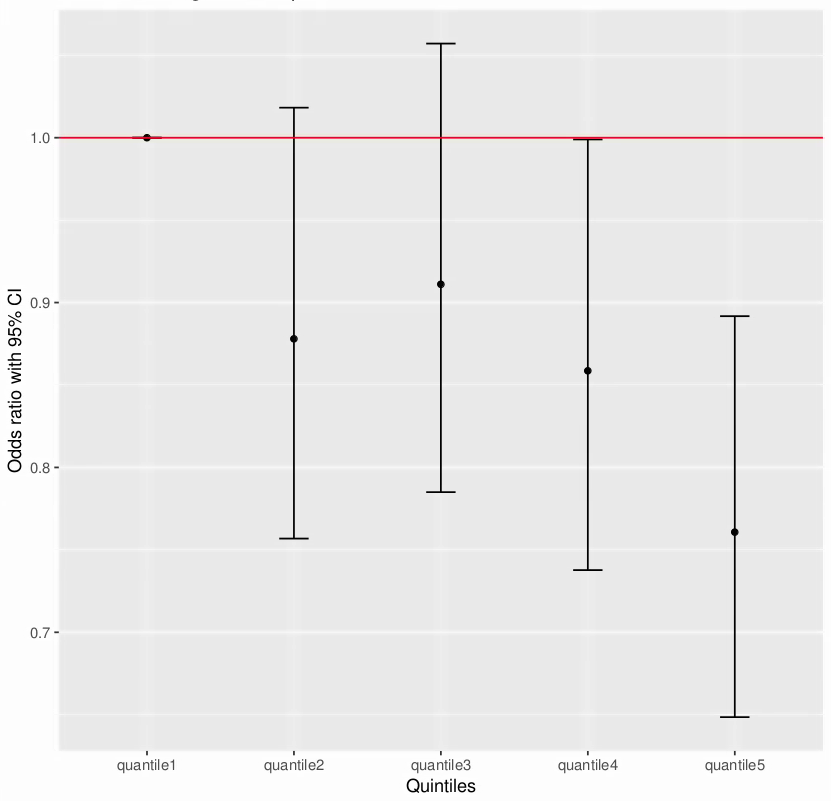

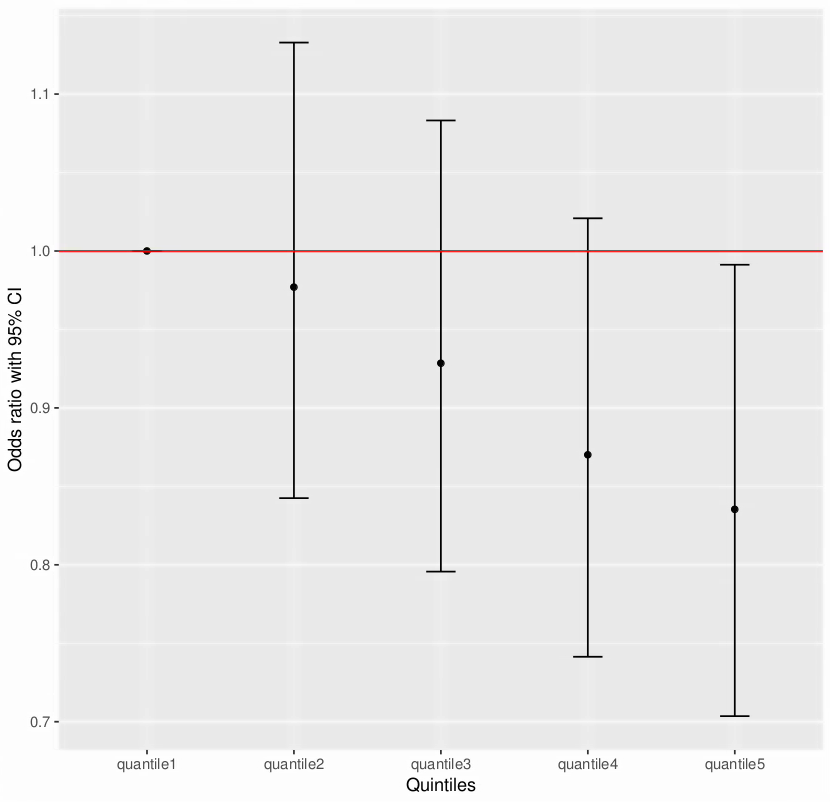

D. OR for ADHD+DBDs compared to ADHD without DBDs by PRS for college completion.

F. OR for ADHD+DBDs compared to ADHD without DBDs by PRS for verbal numerical reasoning.

E. OR for ADHD+DBDs compared to ADHD without DBDs by PRS for human intelligence.

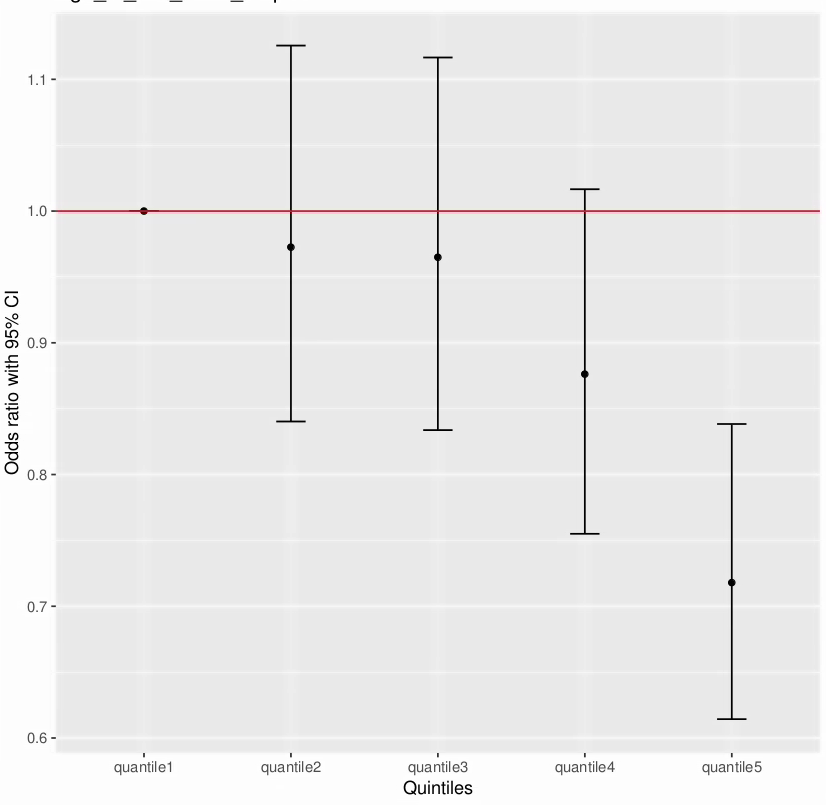

G. OR for ADHD+DBDs compared to ADHD without DBDs by PRS for age at first birth.
